# Gastrodia-Uncaria Water Extract Inhibits Endoplasmic Reticulum Stress and Matrix Metalloproteinase in Protecting against Cerebral Ischemia

**DOI:** 10.1101/2021.07.21.452303

**Authors:** Choi Angus Yiu-Ting, Xian Jia Wen, MA Sum Yi, Lin Zhixiu, Chan Chun Wai

## Abstract

Stroke is the second leading cause of death in worldwide, in which cerebral ischemia accounts for 87% of all cases. The building up of endoplasmic reticulum stress in cerebral ischemia contributes to the disruption of blood brain barrier and neuronal cell death. The only FDA-approved drug, recombinant tissue plasminogen activator, is still of limited use due to the narrow window period and lack of neuroprotective effect. Therefore, it is necessary to explore alternative treatment on cerebral ischemia. *Tianma-Gouteng* decoction is a traditional Chinese Medicine prescription used to treat brain diseases in China. In this study, we investigated the neuroprotective effect of a water extract consisting of *Gastrodia elata* and *Uncaria rhynchophylla*, which are the two main herbs in the decoction. Cerebral ischemia was induced in rats using middle cerebral artery occlusion. GUW-treated rats have significantly reduced infarction volume and recovered neurological functions. The number of protein aggregates and caspase-12 positive cells were significantly inhibited. *In vitro* oxygen-glucose deprivation / reoxygenation stroke model demonstrated that the unfolded protein response proteins GRP78 and PDI were upregulated by GUW. Less ubiquitin puncta and normalized ubiquitin distribution indicated the reduction in endoplasmic reticulum stress. Furthermore, a lower Evan blue signal and MMPsense signal was observed, suggesting that GUW may preserve the blood brain barrier integrity through inhibiting MMP activity. Taken together, this suggested that GUW protected ischemic neurons and the blood brain barrier through inhibiting endoplasmic reticulum stress.

## INTRODUCTION

Stroke is the second leading causes of death in worldwide, in which 87% of the stroke cases were cerebral ischemia [1]. After stroke, more than 60% of patients would experience neurological impairments and loss in locomotor functioning [1]. Recombinant tissue plasminogen activator (tPA) is the only prescribed medicine in worldwide. However, the shortcoming of tPA treatment are the narrow therapeutic window and increasing risk of intracerebral hemorrhage. Therefore, it is a challenge to seek for alternative treatment methods.

Cerebral ischemia is defined by the disruption of blood flow towards the brain tissue. Such a blockage is caused by the formation of thrombus and embolus, a plaque that could obstruct the blood vessel [2]. After the removal of the blood clot through mechanical thrombectomy or by using recombinant tissue plasminogen activator, blood supply is restored and reperfusion phase proceeds. During this phase, the calcium stores in the ER will be depleted, resulting in a decrease in protein folding efficiency and the accumulation of unfolded protein aggregates [3]. Thus, endoplasmic reticulum stress is built up. The unfolded protein response (UPR) is a mechanism to counteract ER stress [4]. Chaperones such as GRP78, protein disulfide isomerase (PDI) are upregulated to assist in proper folding of the protein aggregates to attenuate ER stress [5]. Nonetheless, if ER stress persists for a long period of time, caspase-12 is activated to trigger cell death [6].

The increase in ER stress leads to the activation of matrix metalloproteinases (MMP), causing the disruption of blood brain barrier [7]. The activation of MMP during cerebral ischemia can cause irreversible loss of blood brain barrier (BBB) integrity, edema formation, hemorrhagic transformation, and further neuronal cell death [8,9]. Inhibition of MMP may protect BBB disruption through reducing brain microvascular endothelial cell damage [10]. The above indicated inhibiting ER stress may be beneficial in protecting against BBB dysruption. *Gastrodia elata* (GE) and *Uncaria rhynchophylla* (UR) are the two main herbs in *Tianma-Gouteng* decoction, a Chinese Medicine prescription that is used to treat head related disease such as stroke. We have previously found that GE and UR in the *Gastrodia-*Uncaria water extract (GUW) work synergistically to protect against cerebral ischemia through inhibiting oxidative stress and apoptosis [11]. Metabolomics analysis of the cerebral spinal fluid revealed changes in the amount of endoplasmic reticulum stress-related metabolites in GUW-treated ischemic rats [12]. In this study, we investigated the effects of GUW in protecting against cerebral ischemia-induced apoptosis through reducing ER stress and preserving blood brain barrier integrity.

## METHODS AND MATERIALS

### Reagents

All chemicals, culture medium and secondary antibodies are purchased in Thermo Fisher Scientific (Carlsbad, CA, USA) unless specified. Primary antibodies were purchased from Abcam (Cambridge, UK).

### Herbal extract preparation

GUW, GE, and UR were prepared as previously mentioned [11]. HPLC of the herbal extracts were performed with the chemical markers for quantification and authentication [11].

### Middle cerebral artery occlusion (MCAO)

All animal experimental procedures were conducted with the approval from the Animal Ethics Committee of the Chinese University of Hong Kong (ref. No.: 11/068/MIS-5). MCAO was adopted from previous study [11]. Briefly, male Sprague Dawley rats of 260–280 g were used for the study and were housed in a 12 h: 12 h light-dark cycle environment, fed with chow and water *ad libitum*. The rats were first anesthetized with 400 mg/kg chloral hydrate (VWR International; Poole, England) injected intraperitoneally. The right external carotid artery was excised and ligated. A round-tipped 4-0 nylon suture was inserted towards the middle cerebral artery via the internal carotid artery. The middle cerebral artery was occluded by the nylon suture for 2 h. The suture was then removed and the external carotid artery was cauterized. The operated rats were caged individually and warmed with a heating lamp. The same operation procedures were carried out on the shams rats but without inserting the nylon suture to occlude the middle cerebral artery.

### GUW treatment and assessment

GUW powder was dissolved in ddH_2_O and was orally administered to the MCAO rats with a human equivalent dosage of 288.6 mg/kg GUW once daily for seven consecutive days [13]. One group of rats subjected to MCAO received GUW treatment. The rest of the MCAO rats and sham rats are fed with ddH2O. After treatment, neurological deficit score and triphenyl tetrazolium chloride (TTC) staining were performed to assess the brain infarct volume and neurological functioning as previously described [11].

### Cresyl violet staining

Another set of brains were harvested and fixed in buffered 10% formalin through intracardial perfusion. They were then dehydrated in 70% ethanol. The slices were dehydrated in 100% ethanol and embedded in wax. Brain sections were cut at 5 µm thickness. After hydration of the slides using a decreasing ethanol gradient, the slides were placed into the cresyl violet solution for 10 min at 60°C. The slides were then dehydrated and mounted with DPX. The total number of cells with intensely stained Nissl bodies were counted in five random views in the penumbra region using ApoTome.2 fluorescence microscopy (Carl Zeiss AG; Oberkochen, Germany).

### Evans blue assay

Evans blue staining was performed to evaluate the effect of GUW treatment on BBB permeability after MCAO-induced injury. At day 7 of post-operation, the animals were infused with 2 ml/kg of 2% Evans blue dye in saline (Sigma Aldrich, MO, USA) through the tail vein. After 2 h, rats were anesthetized and transcardially perfused with PBS until a colorless perfusion fluid was obtained from the right atrium. The brains were harvested and washed with PBS. brain imaging was conducted via in vivo MS FX-Pro system (Carestream Health, Inc., NY, USA) at an excitation wavelength of 620 nm and emission wavelength of 700 nm. The near infrared fluorescent signal in region of interest (ROI) of ischemia hemisphere was measured as compared with the contralateral hemisphere of brain [14]. The target-to-background ratio (TBR) of near infrared fluorescent signal was calculated.

### *In vivo* and *ex vivo* near-infrared fluorescence imaging

The near infrared fluorescent dye MMPsense 680 (PerkinElmer Inc, MA, USA) was used to monitor the MMP activity. The MMPSense 680 was injected intravenously through tail vein with a volume of 200 μl, equivalent to 4 nmol per rat at day 2 of post-stroke. After anaesthetized with 1% isoflurane gas by inhalation, the rat is put into in vivo MS FX-Pro system and exposed with an excitation wavelength of 650 nm, the near-infrared fluorescent signal (NIRF) of MMPsense 680 is measured at emission wavelength of 700 nm at day 3, 5 and 7. At day 7 post-operation, the whole brain is harvested. The MMPsense 680 NIRF signal of harvested whole brain and bran slices in 2mm thick was measured. The near infrared fluorescent signal in RIO is measured as compared with the contralateral hemisphere of brain. TBR of near infrared fluorescent signal was calculated as previously mentioned [15].

### Immunohistochemistry

Antigen retrieval was carried out by placing the slides in boiling citrate buffer for 20 min. The tissues were blocked with 10% goat serum, 1% BSA in PBS for 1 h. The slides were then incubated with primary antibodies overnight at 4°C, followed by incubation of secondary antibodies for 1 h at room temperature. The tissues were mounted with ProLong Gold Antifade Reagent with DAPI. The concentrations of the antibody used are as follow: Rabbit anti-Ubiquitin, 1:100; Mouse anti-Caspase-12, 1:100; Goat anti-rabbit IgG (H+L) with Alexa Fluor 488 conjugate, 1:200; Goat anti-mouse IgG (H&L) with Alexa Fluor 594 conjugate, 1:200. Fluorescence microscopy was carried out using ApoTome.2 fluorescence microscopy.

### Cell culture and differentiation

SK-N-SH cells (ATCC HTB-11) were obtained from American Type Culture Collection (ATCC; Manassas, VA). The cells were cultured with RPMI 1640 medium supplemented with 10% FBS and 1% penicillin/streptomycin/neomycin. Cell cultures were incubated under 5% CO_2_ at 80% humidity unless specified. Cells were differentiated with neuronal phenotype used a protocol mentioned previously with modifications in a 35 mm 6-well plate [16]. The plate was pre-coated with poly-L-lysine for neuronal cell adhesion. SK-N-SH cells were then seeded at a density of 1 × 10^5^ cells/well and cultured with RPMI 1640 medium supplemented with 10% FBS, 1% penicillin-streptomycin for the first two days. The wells were washed with FBS-free RMPI 1640 before adding RMPI 1640 supplemented with 1% FBS and 10 µM retinoic acid. The cells were then cultured for four days to obtain the neuronal phenotype.

### Cytoviablity assay of extracts

SK-N-SH cells were seeded in 96-well plates at a density of 15000 cells / well and were differentiated as above mentioned. The culture medium was then replaced with 100 µl of RA containing RPMI 1640 medium added with GUW, GE, and UR of several concentrations. The plate was incubated at 37 °C under air with 21% O_2_ and 5% CO_2_ at 80% humidity for 24hr. After 24 hr, the wells were aspirated and 100 µl of 0.5 mg/ml MTT was added to each well. After incubation for 3 h, the solution was drawn out and 100 µl DMSO was added to each well. An addition lane was added with DMSO to serve as the solvent control. The plate was then placed into a BioTek µQaunt microplate reader (BioTek; Winooski, VT, US) and the optical density at 570 nm was recorded by the Gen5 Data Analysis Software (BioTek; Winooski, VT, US). The cell viability of treated samples was normalized to normoxic control as:

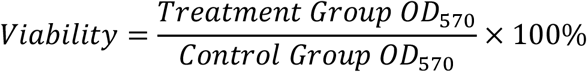

### Oxygen-glucose deprivation / reperfusion (OGD/R)

The induction of OGD was adapted from Tabakman’s method [17]. Neuronal differentiated SK-N-SH in 6-well plates were washed with 1 ml glucose-free RPMI 1640 twice. Afterward, GUW were added at concentrations of either 500 or 1000 µg/ml in glucose-free RPMI 1640 medium. The cell cultures were placed in a hypoxia incubation chamber (Stemcell Technologies; Vancouver, BC, Canada) filled with 95% N_2_, 5% CO_2_ (Hong Kong Specialty Gases Co. Ltd.; HK, China). The cultures in the hypoxia chamber were kept for 7 h to undergo the ischemic phase. After OGD, the culture medium was refreshed with glucose containing RPMI 1640 supplemented with 1% FBS, 10 µM RA and 1% penicillin/streptomycin/neomycin and were keep under normoxic condutions for 17 h to carry out the reoxygenation phase. Normoxic controls were incubated at 37 º C normoxic condition with the differentiation medium at the same time point with the OGD group.

### Immunocytochemistry

The amount of unfolded proteins in terms of ubiquitinated aggregates and neuronal morphology was examined using immunocytochemistry. Glass coverslips (Paul Marienfeld GmbH & Co. KG; Lauda-Königshofen, Germany) were coated with poly-L-lysin and placed in six-well plates. The cells were fixed with 4% paraformaldehyde in PBS for 10 min at room temperature and permeabilized using 0.01% Triton X-100 in PBS. The cells were blocked with 10% goat serum, 1% BSA in PBS-T for 1 hr. The cells were then incubated with primary antibodies in 10% goat serum, 1% BSA in PBST overnight at 4°C, followed by incubated with secondary antibodies in 10% goat serum, 1% BSA in PBST for 1 h at room temperature. The glass coverslips are mounted on glass slides with ProLong Gold Antifade Reagent with DAPI. The concentrations of the antibody used are as follow: Rabbit anti-Ubiquitin, 1:100; Mouse anti-Neurofilament 160kDa, 1:100; Goat anti-rabbit IgG (H+L) with Alexa Fluor 488 conjugate, 1:400; Goat anti-mouse IgG (H+L) with Alexa Fluor 594 conjugate, 1:200. Images were captured using ApoTome.2. Ubiquitinated protein aggregates is defined as the green puncta present in the image. Two neurons are defined as connected when the axons were observed linked together. The number of puncta and connections were counted in five different regions within a sample.

### Statistical analysis

Results obtained were statistically compared using SPSS20 (IBM.; North Castle, NY, USA). One-way ANOVA *post hoc* Tukey’s test was used in all data sets besides from cytotoxicity test and OGD/R cell viability assessment. One-way ANOVA *post hoc* Dunnet’s test was used on the above mentioned two data set. Student’s t-test was used in Evan’s blue and MMPsense assay. *p* < 0.05 was considered to be statistically significant.

## RESULTS

### GUW protected against cerebral ischemia

The neurological deficit score was assessed on day 1 and 7 post-operation (Fig. 1a). At day 1, a significant increase in neurological deficit score was observed in MCAO (*p* < 0.001) and GUW rats (*p* < 0.001) when compared with sham group rats; insignificant difference (*p* > 0.05) was found between MCAO and GUW rats. GUW rats had significant improvements in neurological deficits on day 7 when compared with the score on day 1 (*p* < 0.001) and MCAO rats on day 7 (*p* < 0.001). While the mean neurological deficit score didn’t significantly improve in the MCAO rats at day 7 (*p* > 0.05) and remained significantly higher than the sham rats (*p* < 0.001), insignificant difference was found between GUW rats and sham rats on day 7 (*p* > 0.05). TTC staining was performed to examine the infarction volume (Fig. 1b). The brain was significantly infarcted by 45.6% after MCAO (*p* < 0.001, Fig. 1c). GUW treatment rats, the infarct volume was significantly reduced by 55.7% when compared with MCAO rats (*p* < 0.001).

**FIG. 1.**
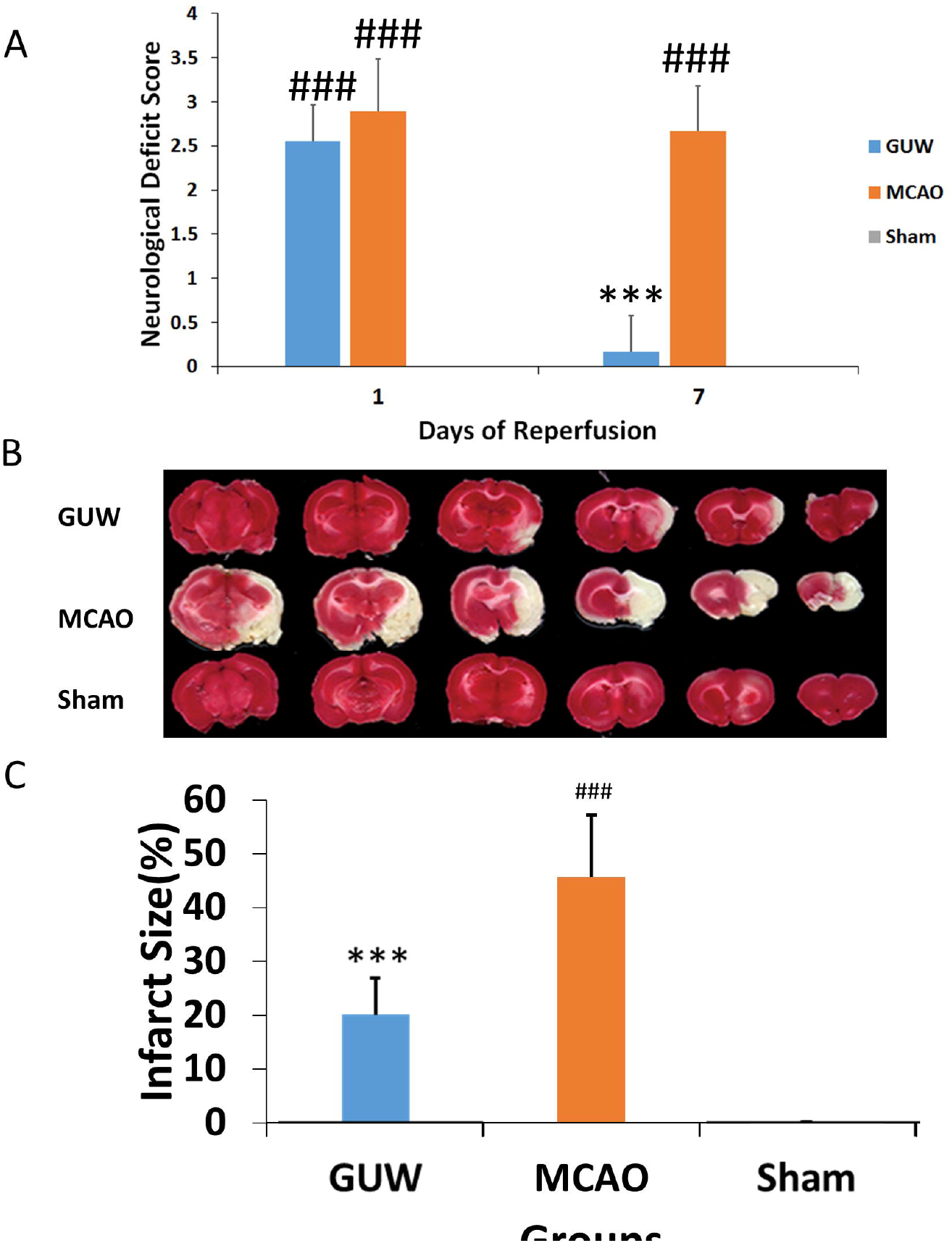
Neurological assessments of rats after MCAO. (A) GUW, MCAO and Sham Rats were subjected to neurological deficit score. (B & C) TTC staining was performed in the brain slices of the above mentioned three groups, the volume of the white infarct region was quantified. Data expressed as mean ± SD. n = 6. One-way ANOVA *post hoc* Tukey’s test. vs Sham, ### *p* < 0.001, vs MCAO, *** *p* < 0.001.

### GUW reduced ER stress

To investigate the effect of MCAO and GUW on neurons, cresyl violet staining was used. Under a higher magnification, two types of staining patterns could be observed (Fig. 2a). The first type of cell with faint color staining and larger cell body indicated living neurons. The second type of cells is intensely stained in purple with a small cell size. After MCAO, 33.8% cells were faintly stained, which was significantly fewer than the sham group rats (*p* < 0.001; Fig. 2b). GUW treatment led to a significant increase in this type of cell population by 53.0% (*p* < 0.001), yet it is significantly fewer than that in the sham group rats (*p* < 0.001).

**FIG. 2.**
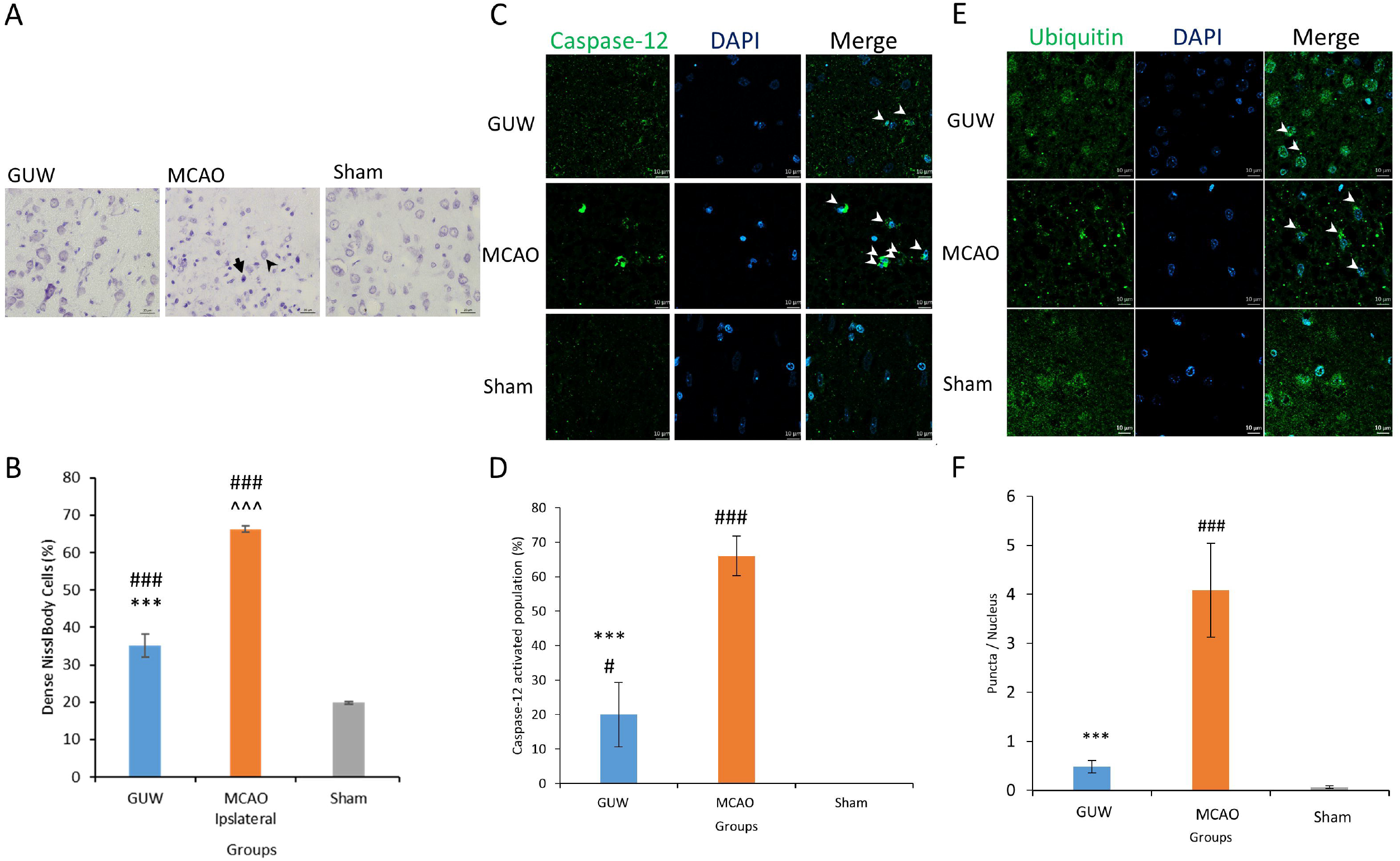
Endoplasmic stress in rats after MCAO. Seven days after MCAO, the brains were harvested to perform cresyl violet staining to assess the number of intensely stained neurons (A). The faintly stained cells were less intensely stained and have a larger cell size (black arrowhead). The intensely stained cells were densely stained and have a smaller cell size (black arrow). The amount of intensely stained cells was quantified (B). Immunohistochemistry of the amount of ubiquitin puncta per cells (C & D) and caspase-12 positive cells in the three groups were quantified (E & F). Data expressed as mean ± SD. n=3. One-way ANOVA *post hoc* Tukey’s test. Vs Sham, ### *p* < 0.001, vs MCAO, *** *p* < 0.001.

Immunohistochemistry was used to study the amount of protein aggregates and protein expression of caspase-12 at the infarct region. In MCAO rats, ubiquitinated protein aggregates were present in the cell body and the ubiquitin distribution was retracted to the soma (Fig. 2c). For GUW group, the number of aggregates was significantly reduced by 88.3% (*p* < 0.001; Fig. 2d). Furthermore, a normal ubiquitin distribution was observed. The caspase-12 expression was upregulated at the infarct region in the MCAO rats when compared to the sham rats (Fig. 2e). The number of caspase-12 positive cells was reduced significantly by 46.6% in GUW-treated rats (*p* < 0.001; Fig. 2f).

### GUW Protected Against OGD/R

The cytotoxicity of GE, UR and GUW were examined using MTT assay (Fig. 3a). After 24 h of incubation, no significant toxicity was observed in GE up to the maximally tested concentration at 4000 µg/ml (99.9%, *p* > 0.05). However, significant toxicity was observed in UR at concentrations above 250 µg/ml (72.74%, *p* < 0.001). Nearly all the cells are non-viable at concentrations above 1000 µg/ml (6.28%, *p* < 0.001). For GUW, insignificant toxicity was observed up to concentrations at 500 µg/ml (95.50%, *p* > 0.05). Based on the cytotoxicity profile, GUW at concentrations of 500 and 1000 µg/ml were used in OGD/R assays. MMT assay showed that GUW at 500 and 1000 µg/ml significantly increased the cell viability by 21.36% and 19.29% after OGD/R (*p* < 0.05; Fig. 3b). GE group showed 13% higher cell viability than the control group (*p* < 0.05). UR concentrations at 50 and 100 µg/ml did not lead to significant protective effect. Furthermore, UR induced significant toxicity when 500 µg/ml UR was administered after OGD/R (*p* < 0.01). The neurite connectivity was observed using immunocytochemistry (Fig. 3c). GUW treatment significantly rescued the connectivity between the neuronal cells by 46.3% after OGD/R insults (*p* < 0.05; Fig. 3d).

**FIG. 3.**
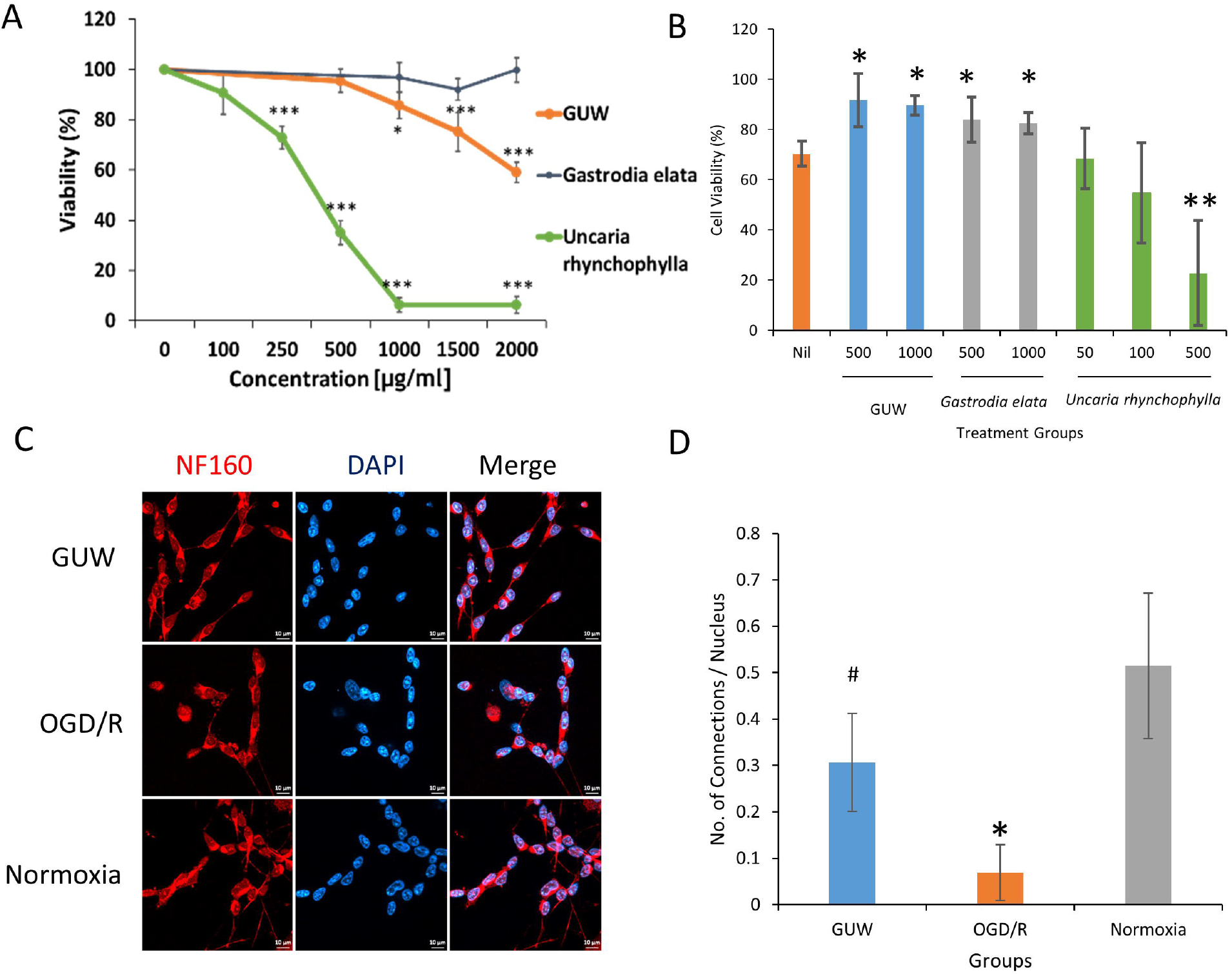
Neuroprotective effect of GUW against OGD/R. (A) The cytotoxicity profile of GE, UR and GUW 24 hr after incubating with differentiated SK-N-SH cells. (B) The cell viability of SK-N-SH cells treated with herbal extracts at various concentrations during OGD/R. (C) Immunocytochemistry using neurofilament 160 kDa (NF160) as a marker to show the neurite formation in SK-N-SH cells after OGD/R. (D) The number of neurite connection per nucleus was evaluated among three groups. Data expressed as mean ± SD. n=3. One-way ANOVA *post hoc* Tukey’s test. Vs Sham, # *p* < 0.05, vs MCAO, * *p* < 0.05.

### GUW protected against ER stress through upregulating UPR

More unfolded proteins were present in the OGD/R neurons (Fig. 4a). Moreover, a retraction of ubiquitin from the neurites to the cell was observed, indicating the occurrence of neurodegeneration. After treating with GUW, the amount of unfolded proteins was reduced significantly by 26.4%, while ubiquitin was normally distributed throughout the neuronal cell (Fig. 4b). Western blot showed that GUW led to an increase in expression of chaperone proteins GRP78, PDI expression (Fig 4c). A downregulation of XBP-1(u) was detected. We further investigated the expression of caspase-12, the initiator caspase of ER stress-induced apoptosis. An upregulation of pro-caspase-12 expression was observed in GUW treatment groups.

**FIG. 4.**
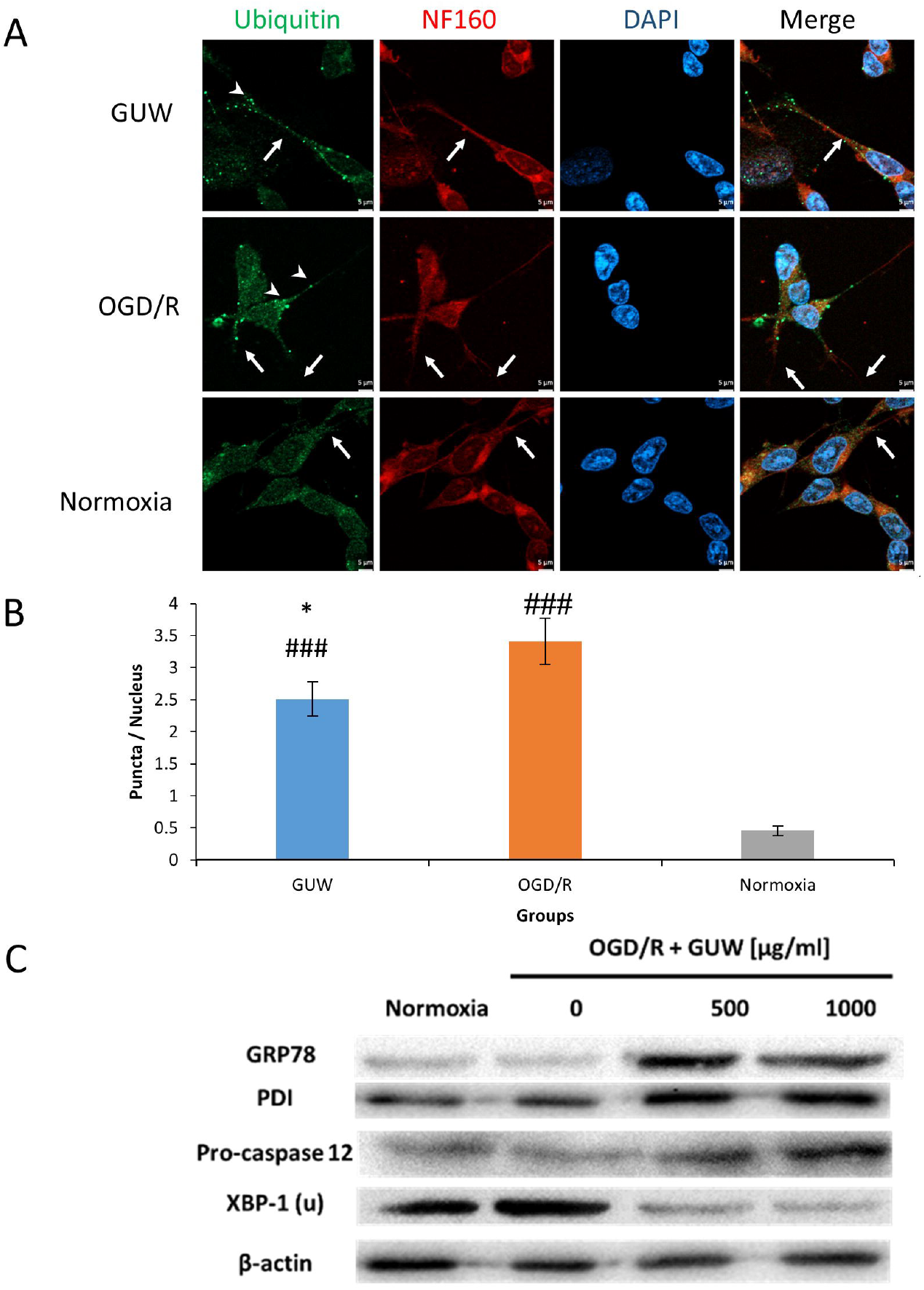
Effect of GUW against endoplasmic reticulum stress. (A) Fluorescence microscopy of ubiquitin (green) and NF160 (red) staining in SK-N-SH cells. White arrows indicate the status of overlapping of ubiquitin and NF160 fluorescence. Arrowheads indicate the presence of ubiquitin puncta. (B)The number of ubiquitin puncta per nucleus was measured among three groups. Data expressed as mean ± SD. n=3. One-way ANOVA *post hoc* Tukey’s test. Vs Sham, ### *p* < 0.001, vs OGD/R, *** *p* < 0.05. Western blot showing the protein expression of GRP78, PDI, XBP-1(u) and pro-caspase-12.

### GUW treatment reduced BBB impairment

Evans blue signal was detected at the left hemisphere after the induction of ischemic insult at both control and GUW treatment group (Fig. 5a). TBR of Evans blue signal in GUW treatment group was 42.3% lower than MCAO group (*p*<0.001; Fig. 5b).

**Fig. 5.**
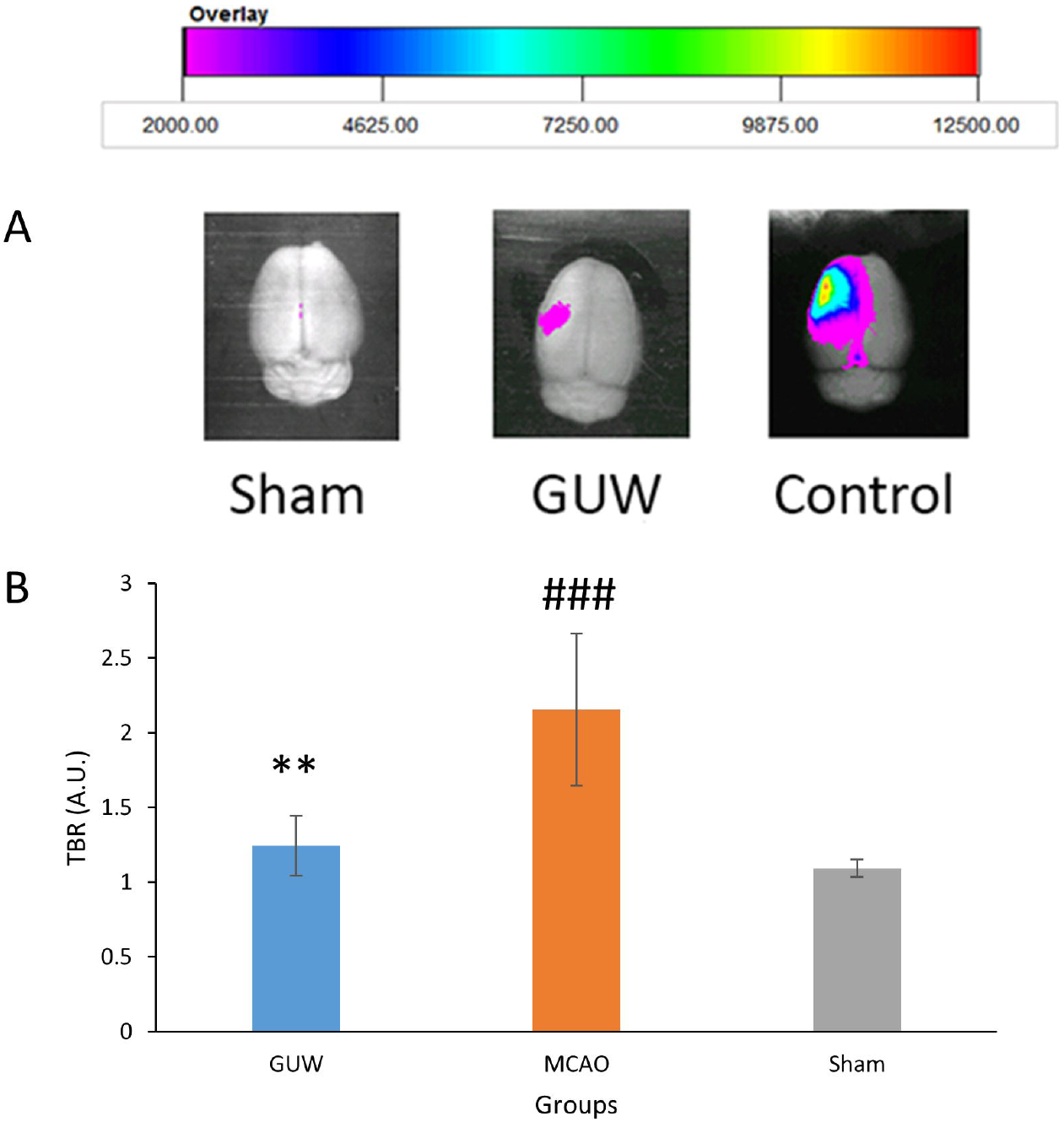
Effect of GUW treatment on Evans blue leakage after ischemia/reperfusion in rats. (A) Representative *ex vivo* NIRF imaging of the rats from different treatment group. At day 7 after MCAO, rats were infused with Evans blue dye and brains were harvested to evaluate the BBB integrity. (B) The Target-to-background ratios (TBRs) calculated from ROI analyses of NIRF image of Evans blue signal. The images were normalized on the color scaling bar. Data are expressed as mean ± SD. n=6. One-way ANOVA *post hoc* Tukey’s test. ^#^ *p* < 0.001 vs Sham group; * *p* < 0.001 vs MCAO.

### GUW inhibited near-infrared fluorescence signal of matrix metalloproteinases activity

NIRF images were obtained at day 3, 5 and 7 of post-operation. A lower fluorescence intensity was observed in the ischemic lesion of GUW treatment group rats compared with MCAO control group (Fig. 6a & 6b). The treatment with GUW led to significant reduction of the fluorescent intensity of matrix metalloproteinases by 25.1 (*p*<0.001), 21.6 (*p*<0.01), and 24.3% (*p*<0.05) at day 3, 5 and 7 respectively (Fig. 6c).

**Fig. 6.**
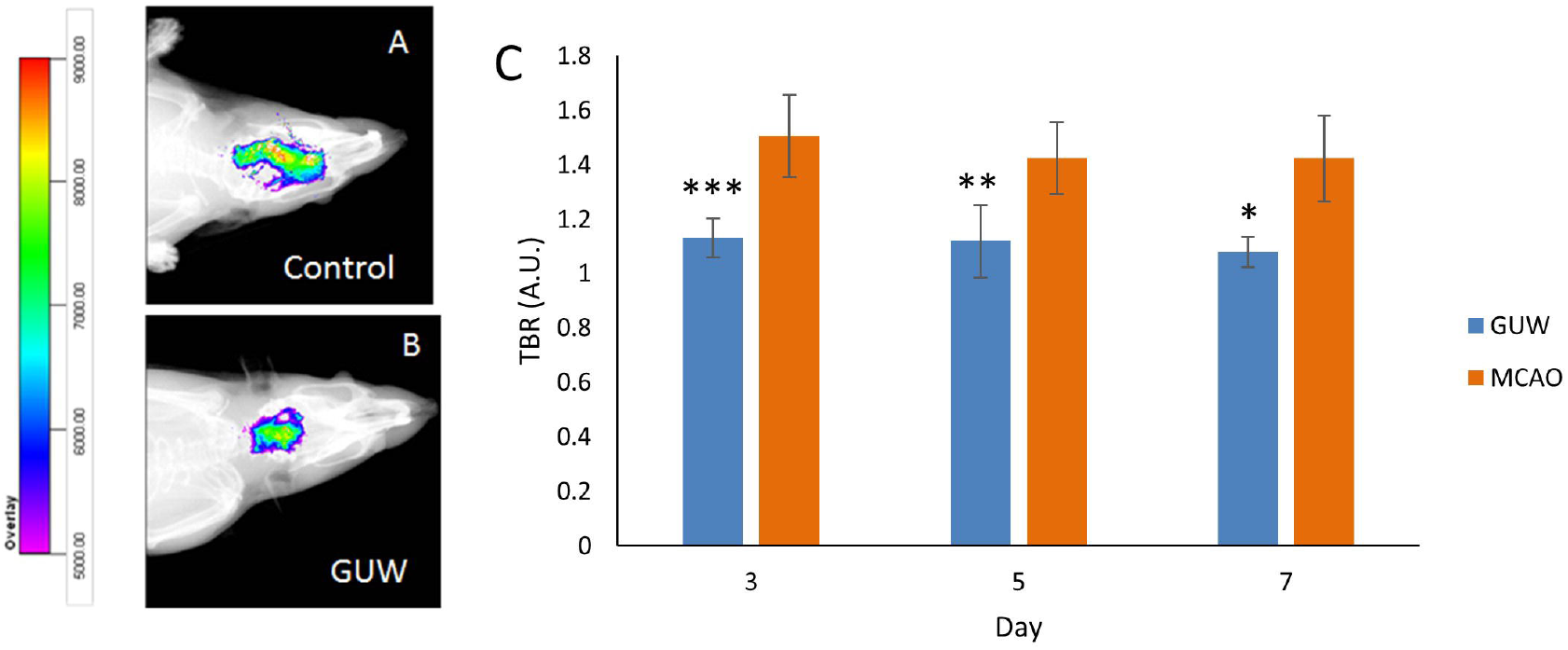
GUW reduced MMP expression. Representative *in vivo* NIRF imaging of the rats at day 5 post MCAO. NIRF imaging was taken after the MMPsense 680 was injected to the brains for 72h. (A) Control group. (B) GUW treatment group. (C) The Target-to-background ratios (TBRs) calculated from ROI analyses of NIRF image. The images were normalized on the color scaling bar. Data are expressed as mean ± SD. Student’s t-test. n=6, * *p* < 0.05, ** *p* < 0.01, \*\*\**p* < 0.001 vs MCAO.

*Ex vivo* brain NIRF imaging was conducted to exclude any possibility of interference caused *in vivo*. At day 7 of post-operation, MMP activities were found in the left hemisphere of MCAO rats (Fig. 7a). The TBR of MMP in MCAO rats was 1.34 fold of sham group (*p*<0.001). GUW treatment led to a significant reduction in the TBR by 26.2% when compared with MCAO rats (Fig. 7c and 7e). NIRF imaging of the brain slices (Fig. 7b and 7d) and from the corresponding brains showed that TBRs in GUW-treated rats were 32.7% lower than that in MCAO group (Fig. 7f; *p* <0.001).

**Fig. 7.**
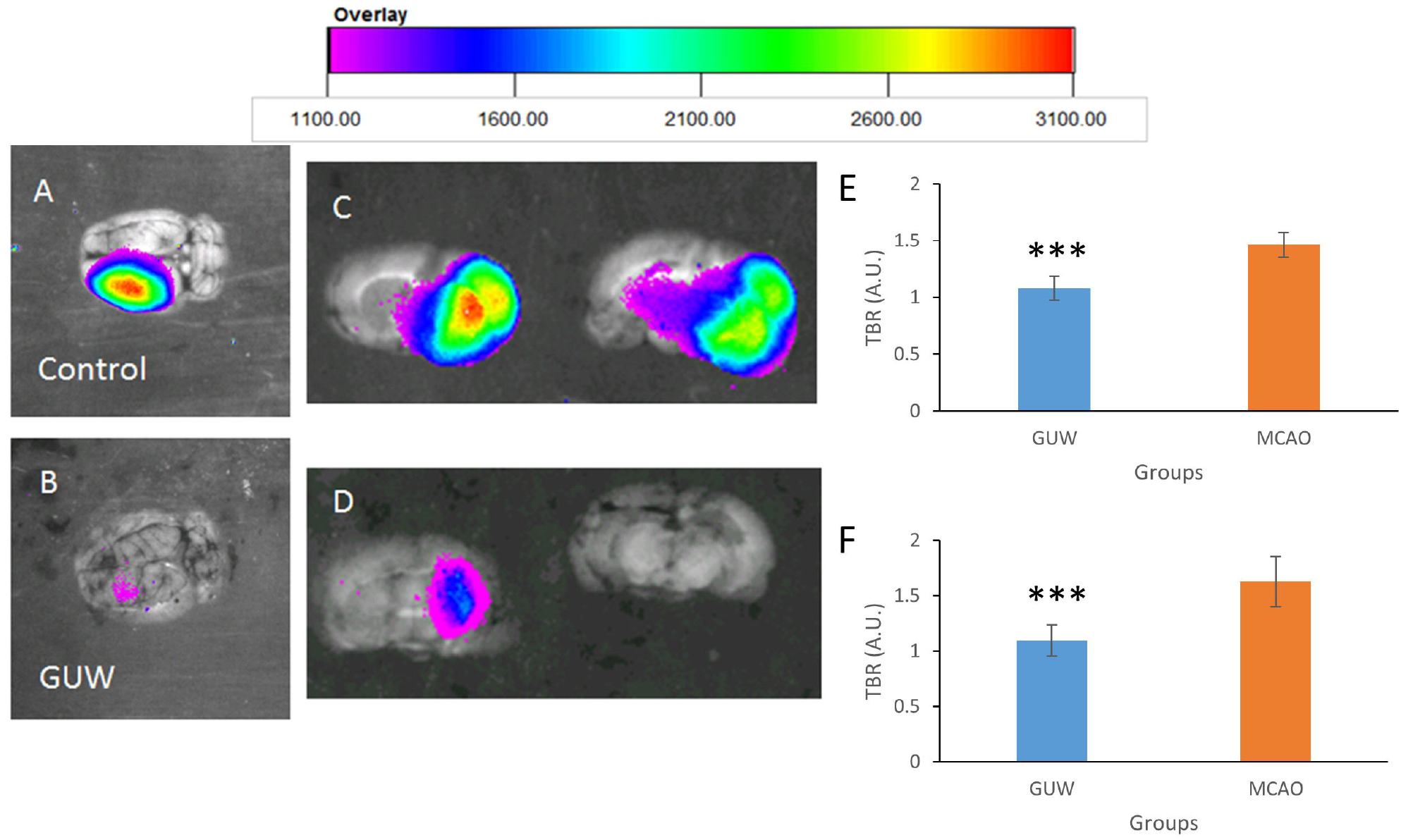
Representative *ex vivo* NIRF imaging of the brains 7 days after MCAO. Brains were harvested at day 7. The MMPsense signal in both the whole brain and 2mm brain slices were captured. (A) Whole brain of control group. (B) Whole brain of GUW treatment group. (C) Brain slices of control group. (D) Brain slices of GUW treatment group. Target-to-background ratios (TBRs) calculated from ROI analyses of NIRF image *ex vivo* of brain. (E) The TBRs of different groups in the whole brain. Data are expressed as mean ± SD. n=6, ^#^ *p* < 0.001 vs Sham group; * *p* < 0.001 vs Control group, by one way ANOVA *post hoc* Tukey’s test. (F) The TBRs of different groups in the brain slices. Data are expressed as mean ± SD. n=6. Student’s t-test. * *p* < 0.001 vs MCAO.

## DISCUSSION

Brain stroke is the second leading causes of death in worldwide. Yet, there is currently no available FDA-approved medication in treating cerebral ischemia at reperfusion phase. In this study, we showed that GUW may significantly protect the brain reperfusion injury followed by ischemic insults through reducing ER stress and preserving BBB integrity.

Seven days of reperfusion was allowed for observing the effect of GUW on recovery. Inignificant difference was found in neurological deficit score between day 1 and day 7 in MCAO rats, which echoed the previous finding that no changes in infarction volume of rat brains were observed after day 1 of reperfusion [18]. Thus, reducing the neurological and locomotor functioning deficits are pivotal in treating stroke. In cortical brain development, the amount of Nissl substance in the small intensely stained neurons positively correlates to the endoplasmic reticulum stress [19]. Cells in MCAO were mostly intensely stained with cresyl violet and small in shape, indicating the loss of cytoplasmic material and the persistence of ER stress. In GUW rats, the amount of intensely stained neurons was much less than that in MCAO rats, though there were still neurons were suffering from ischemic insults. This not only showed that GUW reduced the reperfusion injury on neurons, but also suggested that the endoplasmic reticulum stress in neurons may be inhibited in terms of the reduce in Nissl bodies.

GUW treatment reduced the number of aggregates and caspase-12 after MCAO [20]. This suggested that GUW suppressed ER stress *in vivo* to protect against endoplasmic reticulum stress. Ischemic rats were found to have protein aggregates formed only in the ipsilateral side of the ischemic brain after 30 minutes of reperfusion [21,22]. Proteome analysis revealed 520 proteins being ubiquitinated after cerebral ischemia. Among the protein aggregates, 31% of the proteins within the aggregates are related to protein synthesis and DNA binding, which are essential for cellular survival [23]. As ER stress persists, pro-apoptotic ER stress markers such as caspase-12 and CHOP are found to be significantly upregulated in rat stroke models [24]. In MCAO rat models, upregulation of UPR was found after 7 h of MCAO [24]. Our results demonstrated GUW inhibited ER stress through reducing the accumulation of protein aggregates and thus suppressing caspase-12 activity.

*In vitro* study was carried out to visualize the intracellular events in OGD/R. GE and GUW extract led to a higher SK-N-SH viability. Furthermore, neurodegeneration may occur in CNS neurons after damages, which may lead to neuronal dysfunction [25,26]. After addition of GUW, the dendritic length and connections were maintained as compared with OGD/R group. From western blotting results, it was found that GRP78 and PDI were upregulated by GUW. Chaperones GRP78 and PDI were studied because these two enzymes played important roles in UPR. GRP78 is the master regulator of UPR and it is a major chaperone participating in the protein folding process during UPR [27]. The increase in GRP78 protects neurons against ischemic insults through inhibiting caspase-7 and caspase-12 activity [28,29]. Thus, upregulation of GRP78 is crucial in the protection against reperfusion injury. PDI is a chaperone located in the ER that is responsible for forming disulfide bridges between cysteine residues [30]. Overexpression PDI reduces the number of protein aggregates within motor neurons, suggesting the importance of this chaperone in reducing UPR [31]. Furthermore, PDI could reduce oxidative stress through either reducing disulfide bridges on oxidized substrates or interact with NAPDH oxidase to undergo the reduction reactions [32]. The upregulation of GRP78 and PDI underlined the mechanism of how GUW may reduce the number of protein aggregates to suppress ER stress. Immunocytochemistry could exhibit the aggregates, as the presence of ubiquitin puncta indicates the formation of ubiquitinated-protein aggregates [33]. From our results, there was an increase in the amount of ubiquitin puncta formed within the cell body after OGD/R, while the spots were reduced in GUW. These evidence suggested that GUW might reduce the number of protein aggregates during reperfusion.

The NIRF and Evans blue results suggested that GUW inhibited the expression of MMPs and preserved the BBB integrity. Increase in ER stress may lead to the activation of various MMPs, including MMP-2, MMP-3, and MMP-9 [34,35]. Furthermore, recent studies showed that the upregulation MMPs may enhance ER stress in return and accelerate neurodegeneration [36,37]. In cerebral ischemia, patients with MMP activity at early reperfusion phase is a predictor of better clinical outcome [38,39]. The inhibition effect of GUW on matrix metalloproteinases may play a crucial role in repairing of the ischemic brain, particularly during angiogenesis and reestablishment of cerebral blood flow [40,41].

Together, GUW protected against reperfusion injury through the reduction of endoplasmic reticulum stress by upregulating the chaperones in UPR. Subsequently, the amount of the protein aggregates within the ischemic neurons was reduced and inhibited caspase-12 activation due to ER stress. Therefore, ER stress-induced apoptosis was prevented by GUW treatment in neurons during reperfusion phase.

## ACKNOWLEDGMENTS

This study was supported by the HKSAR Health and Medical Research Fund (Ref: 11120381).

